# RIFT-VAE: grammar-conditioned pretraining and latent-space optimization for RNA inverse folding

**DOI:** 10.64898/2026.08.12.744415

**Authors:** Kentaro Watanabe, Manato Akiyama, Yasubumi Sakakibara

## Abstract

**Background:** RNA inverse folding designs nucleotide sequences expected to adopt a prescribed secondary structure. Search-based solvers can optimize folding-model objectives effectively, but difficult targets can require extensive sampling, and structural optimization alone does not explicitly preserve the sequence distributions or conserved motifs of natural RNA families.

**Methods:** We developed RIFT-VAE, a Transformer-based conditional variational autoencoder that receives a context-free grammar parse-tree representation of a target secondary structure and generates nucleotide labels on the corresponding tree. The framework combines progressively richer grammar rules, self-refinement learning from generated structure-sequence pairs, and cross-entropy-method optimization in the learned latent space. We evaluated RNAfold minimum-free-energy agreement on an RNAcentral-derived test set and the EteRNA100 benchmark, compared the method with four search-based solvers under matched total time budgets, and examined GC-content control and covariance-model family annotation.

**Results:** The complete pipeline achieved RNAfold-Correct/RNAfold-MCC values of 0.833/0.994 on the RNAcentral-derived test set and 0.760/0.977 on EteRNA100. Latent-space optimization accounted for the largest increase in exact structural recovery. Under a 3,600-s total budget on EteRNA100, sequences generated by RIFT-VAE improved the exact-match rate of every tested downstream search method when used as warm starts; the largest change was observed for RNAInverse (Correct, 0.297 to 0.803; MCC, 0.505 to 0.985). The pretrained model also produced sequences with measurable correct-family covariance-model hits and supported explicit GC-content conditioning.

**Conclusions:** RIFT-VAE is best interpreted as a hybrid generative-search framework: pretraining supplies a structure- and family-informed proposal distribution, whereas latent optimization concentrates evaluations in high-scoring regions. The reported structural scores are specific to RNAfold minimum-free-energy validation and do not establish biochemical function. Orthogonal folding predictors, stricter homology-controlled splits, diversity-aware evaluation, architecture-matched dot-bracket ablations, and experimental assays remain priorities for validation.

## Introduction

Recent advances in deep learning have transformed biomolecular structure prediction, including protein structure prediction and the modeling of biomolecular complexes [1, 2]. In parallel, sequence design has emerged as the inverse problem: rather than predicting a structure from a sequence, the objective is to identify sequences compatible with a desired structural or functional specification. Generative protein-design systems have demonstrated the value of learned sequence distributions for proposing candidates compatible with target backbones [3]. Related approaches are now being developed for RNA, including methods conditioned on tertiary structure or tertiary-contact information [4, 11, 12].

RNA inverse folding seeks nucleotide sequences whose predicted secondary structures match a prescribed target. Classical tools such as RNAInverse, MCTS-RNA, SAMFEO, and DesiRNA explore sequence space through local mutation, tree search, multifrontier optimization, Monte Carlo sampling, or replica exchange [5–10]. These methods can obtain high structural accuracy, but the required search can grow with sequence length, structural complexity, and the number of additional design constraints. A trained generative model offers an amortized proposal mechanism: after training, it can sample multiple candidates conditioned on a target and can provide informed initial solutions to downstream combinatorial search.

Structural agreement is nevertheless only one component of functional RNA design. RNA sequences function in cellular environments while interacting with proteins, nucleic acids, ions, metabolites, and other biomolecules. Many families therefore impose sequence-level requirements in addition to a fold. GC content can affect stability and may correlate with function in temperature-responsive RNAs, whereas catalytic, ligand-binding, or recognition motifs may be strongly conserved. In hammerhead ribozymes, for example, conserved catalytic-core and cleavage-site nucleotides are essential for self-cleavage, and preserving these features has enabled functional computational designs [13]. A useful inverse-folding framework should therefore support sequence-level conditions and should retain, where appropriate, family-specific signatures learned from naturally occurring RNAs.

The representation of the target structure is a central modeling choice. A dot-bracket string is compact and widely used, but its serialization presents a nested structure as a flat character sequence. The two nucleotides in a base pair may be far apart in that sequence even though they form a single structural relation. Context-free grammars (CFGs) provide a complementary view: recursive production-rule applications yield a parse tree that directly expresses the nesting of stems and loops and can associate paired positions within the same structural operation. Such a representation provides an explicit interface between classical formal descriptions of RNA secondary structure and sequence-to-sequence neural architectures.

Here, we present RIFT-VAE, a pretrained Transformer-based conditional variational autoencoder (CVAE) for RNA inverse folding. A target secondary structure is converted into a CFG parse tree, serialized as a production-rule token sequence, and augmented with tree-position information. The decoder assigns single-nucleotide or paired-nucleotide labels to terminal-generating nodes, and the tree-labeled output is reconstructed into a 5′-to-3′ sequence. Four components are introduced cumulatively: a conditional latent-variable model, grammar expansion that distinguishes local loop and stem contexts, self-refinement learning (SRL), and cross-entropy-method optimization in latent space.

The study addresses three practical questions. First, can a grammar-conditioned pretrained model generate sequences with high agreement to target secondary structures? Second, can its candidates improve established search-based solvers as warm starts under a matched total computation budget? Third, can the same conditioning interface retain family-level sequence signatures and support GC-content control? We answer these questions on an RNAcentral-derived dataset and EteRNA100, while deliberately limiting our claims to the evaluation actually performed.

## Materials and methods

### Overview and scope of evaluation

Figure 1 summarizes the framework. A target pseudoknot-free secondary structure is supplied in dot-bracket notation and converted into a CFG parse tree. Each production-rule application becomes a structural token, and tree-position encodings are attached to preserve the serialized tree geometry. A Transformer-based CVAE then generates nucleotide labels aligned to terminal-generating nodes. These labels are read in their original 5′-to-3′ order to recover a linear RNA sequence.

**Figure 1.**
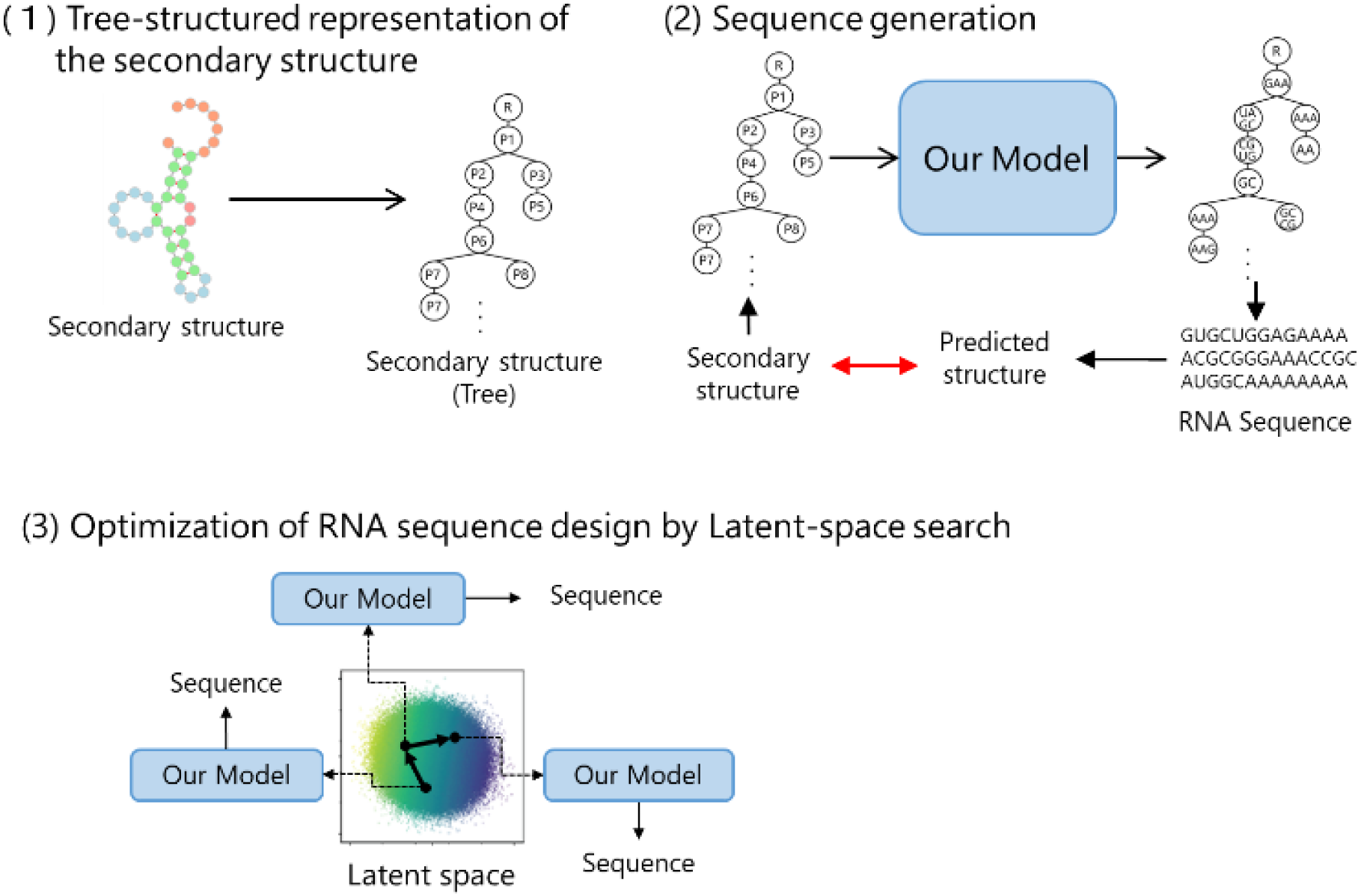
Overview of RIFT-VAE. A target dot-bracket secondary structure is parsed into a CFG tree and serialized as production-rule tokens with tree-position encodings. The grammar-conditioned CVAE generates nucleotide labels on the tree, which are reconstructed into a linear sequence. Candidate sequences are evaluated by RNAfold MFE prediction, and latent-space optimization concentrates sampling in high-scoring regions. The red bidirectional arrow denotes comparison between the target and predicted structures.

Candidate sequences are folded with RNAfold, and the predicted MFE (minimum-free-energy) structure is compared with the target. The same folding model is used for the RNAcentral-derived test set and EteRNA100 to maintain comparability with established inverse-folding evaluations. We use the terms RNAfold-Correct and RNAfold-MCC (Matthews correlation coefficient) when emphasizing that these metrics are predictor-specific. The framework can be used in three modes: direct conditional sampling, latent-space optimization, or generation followed by a search-based solver initialized with the best generated sequence.

### CFG-based tree representation of secondary structure

#### From dot-bracket notation to a typed parse tree

Input secondary structures are provided as dot-bracket strings, where unpaired positions are denoted by ’.’ and paired positions by matching ’(’ and ’)’ characters. We formalize these strings with a CFG and use the corresponding parse tree as the conditioning representation. In the Normal grammar, a structural segment is recursively expanded into unpaired terminals, paired regions, and concatenated subsegments. A simplified example is S -> U S | P S | epsilon, U -> ’.’, and P -> ’(’ S ’)’, where S denotes a structural segment, U an unpaired position, and P a paired region. A dot-bracket string such as ’((..))’ is therefore represented by nested P nodes whose innermost S expands to two U nodes.

The parse tree is traversed in preorder, and each production-rule application is converted into a discrete token. The model input is the resulting production-rule token sequence. To preserve tree geometry after serialization, each token receives an additional tree-position encoding derived from its depth and sibling order. Thus, two identical production rules applied in different structural contexts can be distinguished by their tree positions.

#### Grammar variants and representational granularity

We evaluated four nested grammar designs: Normal, Loop, Loop-Stem, and Loop-Stem α (Figure 2). The Normal grammar expresses the recursive composition of paired and unpaired regions. The Loop grammar adds loop-type-specific nonterminals. The Loop-Stem grammar further distinguishes stem context, and Loop-Stem α increases the amount of local terminal and base-pair-stacking information emitted by one production-rule application. Thus, later variants make differences such as hairpin boundaries, internal-loop asymmetry, stacked pairs, and multibranch junctions more explicit in the local parse-tree topology.

**Figure 2.**
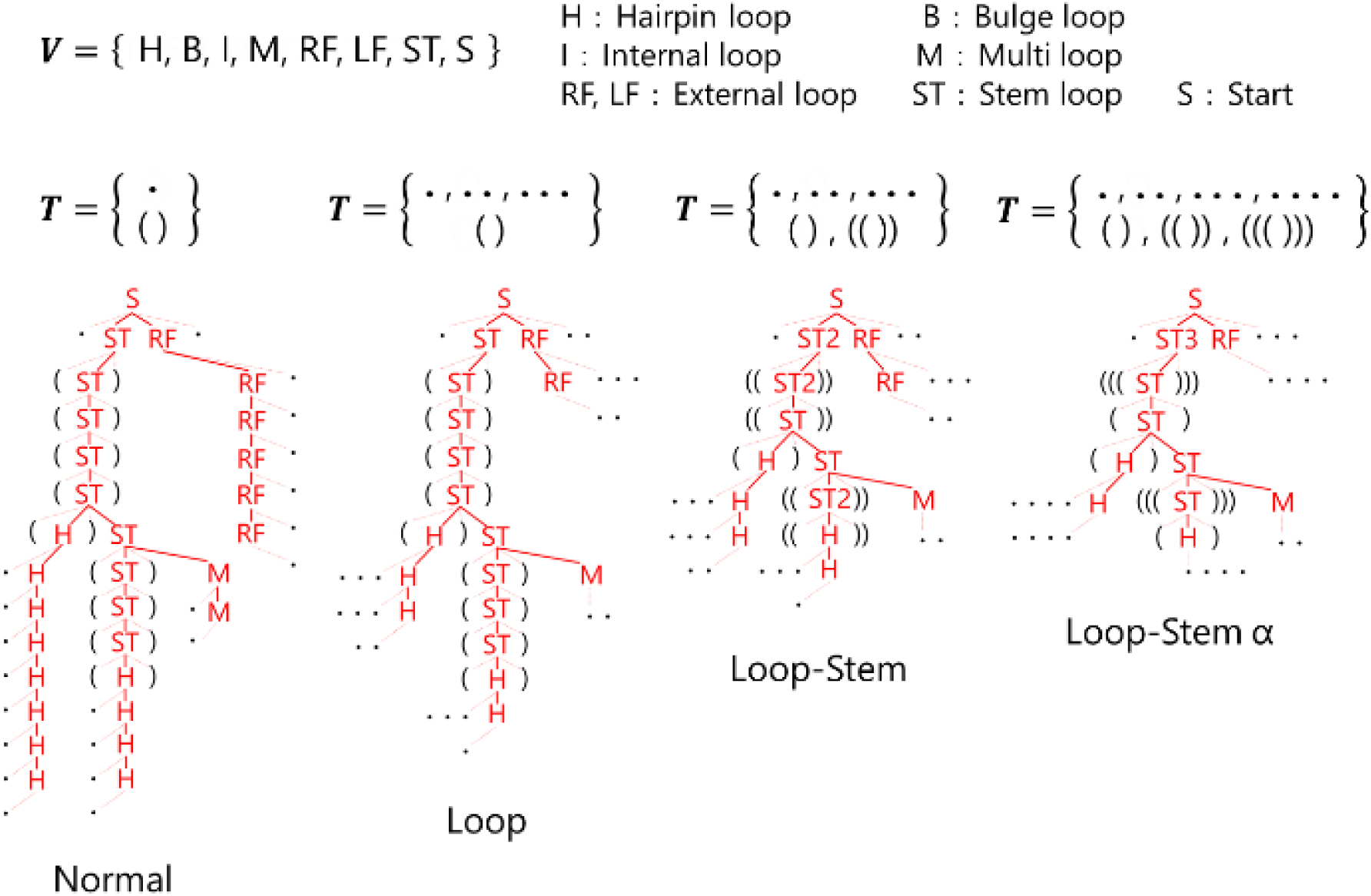
Grammar variants used to represent RNA secondary structure. Nonterminal symbols denote hairpin (H), bulge (B), internal (I), multibranch (M), external left/right (LF/RF), stem (ST), and start (S) contexts. Moving from Normal to Loop, Loop-Stem, and Loop-Stem α increases local terminal-generation granularity and makes loop-stem boundaries, stacked pairs, and branching patterns more explicit in the parse tree.

### Model architecture

#### Base Transformer

The base model is a Transformer encoder-decoder [15] (Figure 3). The encoder receives production-rule tokens together with tree-position encodings. The decoder predicts labels associated with terminal-generating nodes. An unpaired terminal receives one of the four nucleotide labels, whereas a paired terminal receives an ordered nucleotide-pair label. Control and padding symbols are included in the decoder vocabulary. The tree-structured assignment is converted to a linear sequence by reading terminal positions in the original sequence order.

**Figure 3.**
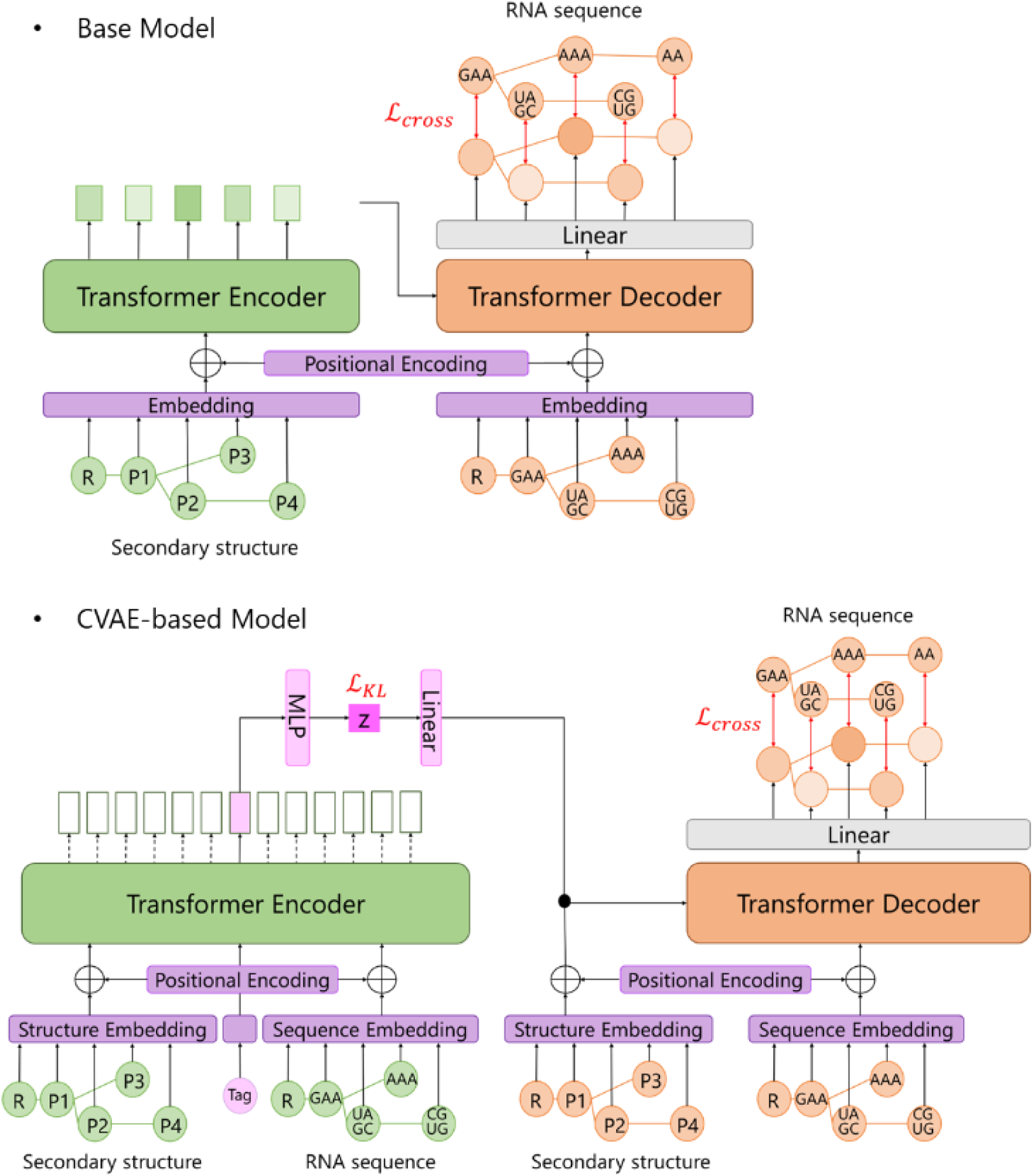
Base Transformer and CVAE architectures. Production-rule tokens and learned tree-position encodings condition the Transformer. The base decoder predicts single- or paired-nucleotide labels on terminal-generating nodes. In the CVAE, the recognition network estimates a latent distribution from the structure-sequence pair during training, and the generative decoder reconstructs nucleotide labels conditioned on the target structure and latent variable z.

The released configuration uses a model dimension of 512, six Transformer layers, eight attention heads, a feed-forward dimension of 2,048, dropout of 0.1, a structural-token vocabulary of 351, a decoder-label vocabulary of 559, and a maximum serialized tree length of 178 nodes. The decoder embedding and output projection share weights.

During training, the target secondary structure is converted into a parse tree and provided as input to the model. The model output, a tree-structured nucleotide sequence in which each terminal node is assigned a nucleotide label, is compared with the true nucleotide labels of the training RNA sequence. The reconstruction loss is the negative log-likelihood of the correct nucleotide labels.

#### Conditional variational autoencoder

RNA inverse folding is one-to-many: multiple sequences may be compatible with one secondary structure. To model this conditional distribution, the deterministic model is extended to a CVAE [16]. Let c denote the grammar-based structural condition, x the nucleotide labels, and z a latent variable. During training, a recognition network approximates qφ(z | x, c), and the decoder models pθ(x | z, c). At generation time, z is sampled from a standard normal prior or from a Gaussian distribution updated during latent-space optimization.

The training objective is the negative evidence lower bound:

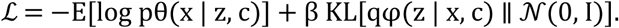

The first term is the nucleotide-label reconstruction loss and the second is the Kullback-Leibler (KL) regularization term. The latent dimension is 128. The initial CVAE configuration is trained with AdamW at a learning rate of 2 × 10⁻⁴, weight decay 0.01, batch size 64, and up to 200 epochs on six GPUs. The public training schedule starts beta at 0.001, increases it to 0.003 after 20 epochs, and to 0.004 after 35 epochs.

For GC-controlled generation, the desired GC fraction is embedded as an additional scalar condition and supplied to the decoder together with the structural condition. This design allows the model to condition simultaneously on a target secondary structure and a sequence-level property without changing the grammar representation.

### Candidate generation and self-refinement learning (SRL)

For the deterministic Transformer, multiple candidates are generated with beam search. For the CVAE, latent vectors are sampled from N(0, I), and each vector is decoded under the target structural condition. We generated 10 candidates per RNAcentral target and 20 candidates per EteRNA100 target and report the best candidate according to the benchmark objective. Best-of-sample reporting evaluates design success but should not be interpreted as a complete characterization of the sampled distribution.

Self-refinement learning (SRL) reuses generated sequences that are inconsistent with their original target under RNAfold. For a generated sequence x conditioned on target c, RNAfold produces a predicted MFE structure ĉ. When ĉ differs from c, the pair (ĉ, x) is treated as a valid positive example because x is, by construction, compatible with ĉ under the same folding model. This converts a failed target-specific generation into additional structure-sequence supervision instead of discarding it.

### Latent-space optimization

We optimize latent variables with the cross-entropy method (CEM) [17, 18]. Starting from a diagonal Gaussian over z, the algorithm samples latent vectors, decodes sequences, predicts their MFE structures with RNAfold, and scores structural agreement. The highest-scoring elite fraction is used to update the Gaussian mean and standard deviation. Repeating this procedure concentrates sampling in latent regions associated with high RNAfold-MCC values.

For EteRNA100, the released configuration permits up to 1,000 CEM iterations, samples 2,048 latent vectors per iteration, retains the top 10% as elites, initializes each latent standard deviation at 1.0, and constrains updated standard deviations to 0.15-3.0. Search stops when an exact solution is found or the time limit is reached. RNAfold evaluations are parallelized across 32 workers, and the ten best unique candidates are retained. RNAcentral uses the same optimization principle under a shorter per-target budget.

Latent-space optimization was run for up to 10 s (seconds) per RNAcentral target and 3,600 s per EteRNA100 target. The latter is the 60-minute computation time conventionally used in prior EteRNA100 comparisons. Direct neural decoding is inexpensive relative to this procedure, but the full optimized pipeline is dominated by repeated folding-model evaluations. Accordingly, we describe RIFT-VAE as a hybrid generative-search framework rather than as an unconditionally faster replacement for search-based solvers.

### Search-based baselines and warm-start initialization

We compared RIFT-VAE with four established search-based methods. RNAInverse applies local sequence mutations to reduce disagreement with the target [5, 6]. MCTS-RNA combines local design operations with Monte Carlo tree search [7]. SAMFEO maintains multiple candidate frontiers and performs structure-aware updates [8]. DesiRNA uses replica-exchange Monte Carlo to improve mixing across complex sequence-folding landscapes [9].

Search-only baselines used the same respective total budgets as latent-space optimization: 10 s per RNAcentral target and 3,600 s per EteRNA100 target. In the EteRNA100 warm-start experiment, RIFT-VAE used 1,800 s for latent-space optimization, and the best candidate initialized a downstream solver that ran for an additional 1,800 s. Thus, the complete warm-start pipeline and each search-only baseline received 3,600 s.

### Datasets and benchmark construction

RNA sequences were obtained from RNAcentral [14]. We retained sequences of at most 500 nucleotides and focused on families with characteristic secondary structures, including rRNA, tRNA, snRNA, snoRNA, miRNA, RNase P RNA, hammerhead ribozymes, other ribozymes, Y RNA, and RsmA/MRP RNA (Figure 4). For every sequence, RNAfold was run under its default MFE settings to obtain a pseudoknot-free dot-bracket target.

**Figure 4.**
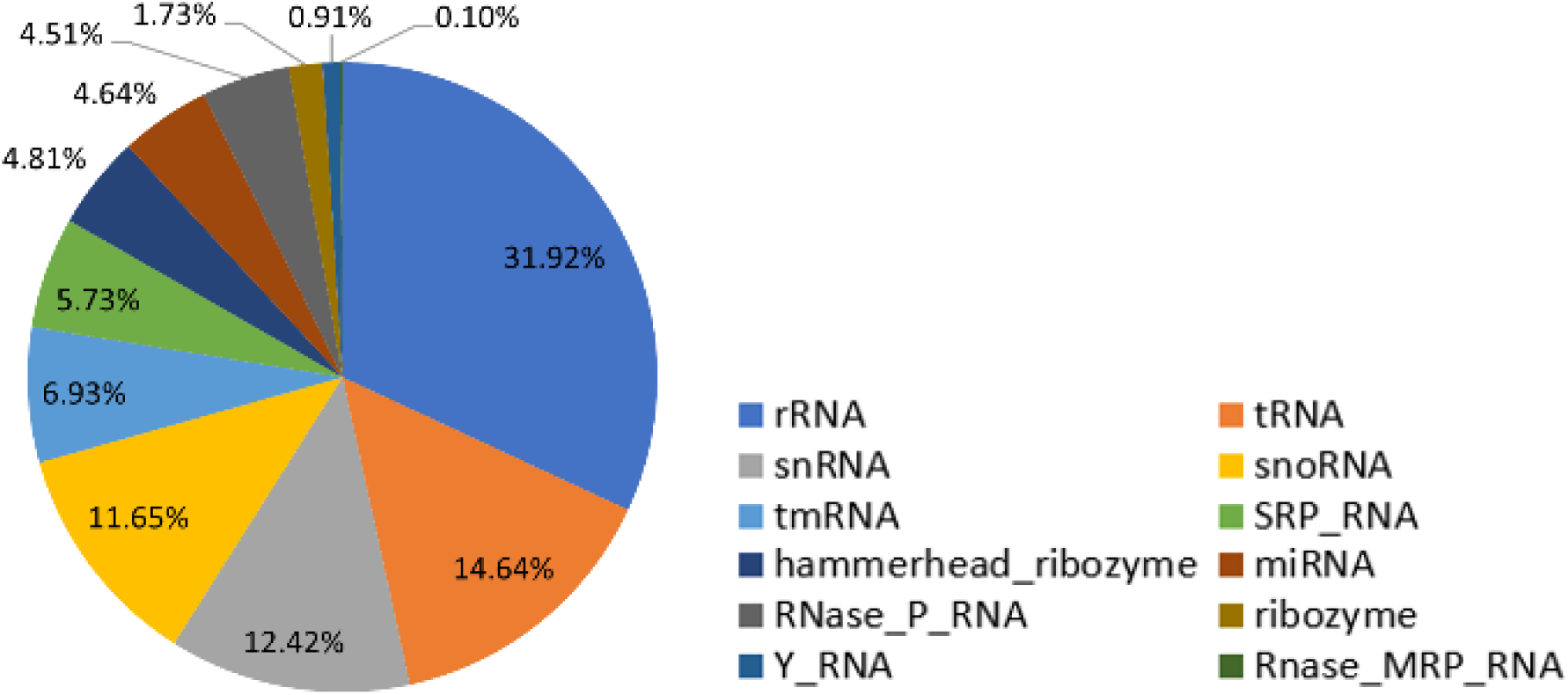
RNA family composition of the RNAcentral-derived dataset. The distribution is imbalanced across families, and aggregate metrics are therefore influenced by the largest categories. Family-level results should be interpreted together with this distribution.

The RNAcentral-derived dataset contains 2,128,591 training examples and 43,118 test examples. The split removes exact duplicate RNAfold-predicted secondary-structure strings across the training and test partitions and removes duplicate target structures within the test set. This criterion prevents exact structural duplicates but does not remove all homologous sequences, near-duplicate folds, remote family relationships, or Rfam clan-level redundancy. The family distribution is markedly imbalanced, and the largest families can dominate aggregate performance.

For a family-level holdout analysis, scaRNA, scRNA, and telomerase RNA were excluded from training. Because distantly related families or structurally similar members of the same clan may remain in the training data, this experiment is described as a family-label holdout rather than strict evolutionary generalization.

EteRNA100v2 contains 100 secondary-structure design targets spanning a range of difficulties [10]. As in prior inverse-folding evaluations, a candidate is successful when RNAfold predicts the specified target as its MFE structure. This benchmark measures compatibility with the ViennaRNA folding engine, not agreement with an experimentally determined structure.

### Evaluation metrics

RNAfold-Correct is 1 when the RNAfold-predicted MFE structure of a designed sequence exactly equals the target and 0 otherwise. Reported Correct values are the fraction of targets solved exactly. RNAfold-MCC is the Matthews correlation coefficient between the upper-triangular base-pair adjacency matrices of the predicted and target structures. With TP, FP, TN, and FN defined over all possible nucleotide pairs, MCC is:

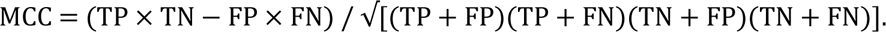

The correct-family annotation rate was evaluated with Infernal 1.1.5 cmscan [19]. A generated sequence was counted as correctly annotated when it matched the covariance model of the target family above the family-specific gathering (GA) cutoff. This metric detects compatibility with known family sequence-structure signatures; it is not a direct functional assay, and lack of a hit does not prove non-functionality.

GC-content control was evaluated using the signed difference between the generated and target GC fractions. We additionally used the bounded match score 1 − |g − t|, where g and t denote the generated and target GC fractions, respectively. Higher values indicate closer agreement with the specified condition.

For EteRNA100, reported RIFT-VAE values are averages over three independent runs. The code, preprocessing pipeline, grammar definitions, pretrained models, and data resources required to reproduce the experiments are available through the repositories listed in the Data availability and Code availability statements.

## Results

### Progressive improvements from four components

Tables 1 and 2 report a cumulative progression from the deterministic Transformer to the full pipeline. On the RNAcentral-derived test set, the base Transformer achieved RNAfold-Correct/RNAfold-MCC values of 0.312/0.815. Introducing the CVAE increased these values to 0.468/0.944. Expanding the grammar to Loop-Stem α produced a further increase to 0.499/0.965, and SRL raised performance to 0.654/0.978. Latent-space optimization yielded the largest final improvement, reaching 0.833/0.994.

**Table 1.** Cumulative performance progression on the RNAcentral-derived test set. Each row adds one component to the preceding configuration.

| **Model** | **RNAfold-Correct** | **RNAfold-MCC** |
| --- | --- | --- |
| Base Transformer | 0.312 | 0.815 |
| + CVAE | 0.468 | 0.944 |
| + Grammar expansion | 0.499 | 0.965 |
| + SRL | 0.654 | 0.978 |
| + Latent-space optimization | 0.833 | 0.994 |

**Table 2.**
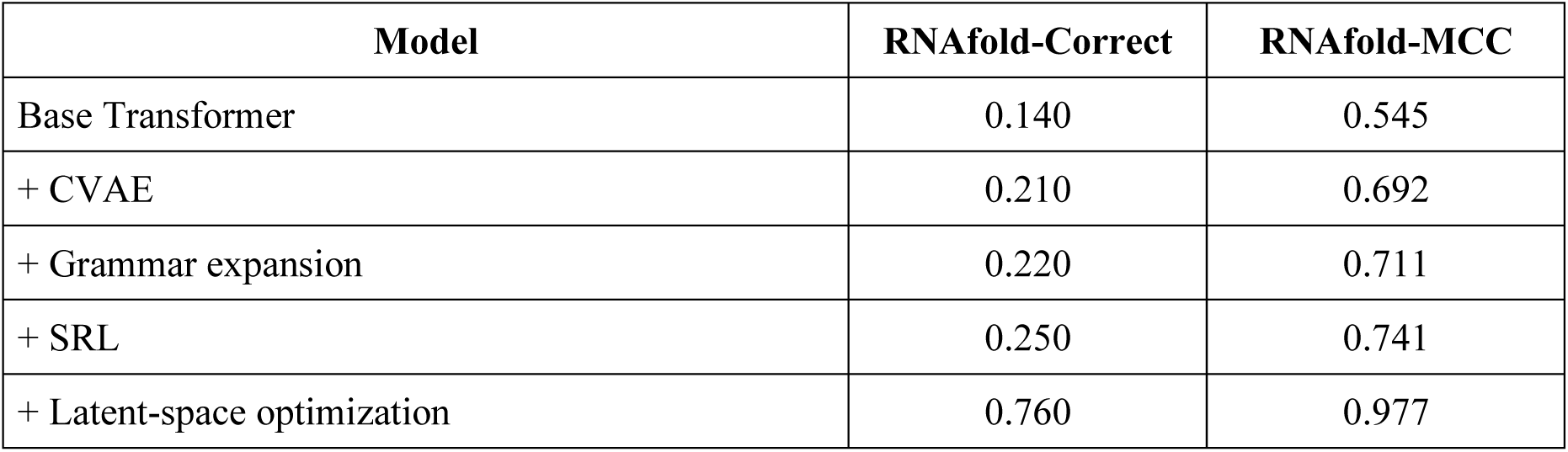
Cumulative performance progression on the EteRNA100 benchmark. Values are averages over three independent runs.

| **Model** | **RNAfold-Correct** | **RNAfold-MCC** |
| --- | --- | --- |
| Base Transformer | 0.140 | 0.545 |
| + CVAE | 0.210 | 0.692 |
| + Grammar expansion | 0.220 | 0.711 |
| + SRL | 0.250 | 0.741 |
| + Latent-space optimization | 0.760 | 0.977 |

The same pattern was observed on EteRNA100. The base Transformer achieved 0.140/0.545, the CVAE 0.210/0.692, grammar expansion 0.220/0.711, and SRL 0.250/0.741. CEM optimization increased the result to 0.760/0.977. Thus, each cumulative component improved both metrics, but most exact-match recovery in the final EteRNA100 pipeline came from latent-space search rather than one-shot generation.

### Family-label holdout analysis

RIFT-VAE maintained RNAfold-MCC values near 1.0 and Correct values comparable to those of included families for scaRNA, scRNA, and telomerase RNA, which were excluded from training by family label (Figure 5). This observation indicates that the model can generate high RNAfold-agreement candidates for target structures assigned to these held-out labels.

**Figure 5.**
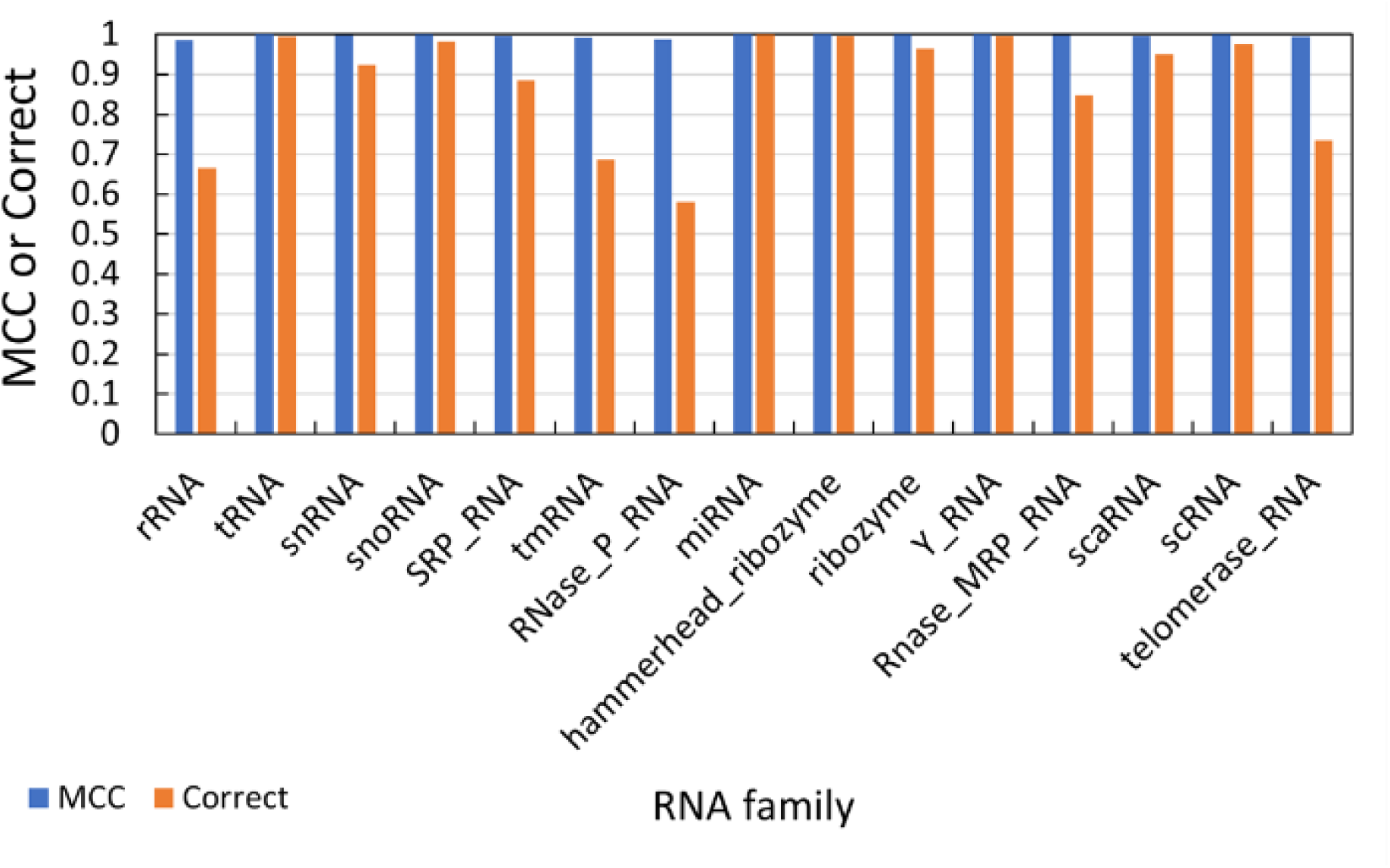
Per-family RNAfold-MCC and RNAfold-Correct values. scaRNA, scRNA, and telomerase RNA were excluded from training by family label. The analysis is a family-label holdout, not a clan-level or homology-controlled generalization test.

### EteRNA100 benchmark and warm-start utility

Under the full 3,600-s setting, RIFT-VAE achieved RNAfold-Correct/RNAfold-MCC values of 0.760/0.977 on EteRNA100 (Table 3). Its exact-match rate was similar to SAMFEO (0.770) and lower than DesiRNA (0.907). DesiRNA was also strongest in MCC (0.996). These results position RIFT-VAE as competitive with several established solvers but not as the best method when RNAfold exact-match success is the sole objective.

**Table 3.** EteRNA100 comparison with search-based methods and warm-start hybrids under a matched total time budget.

| Model | RNAfold-Correct | RNAfold-MCC |
| --- | --- | --- |
| RIFT-VAE | 0.760 | 0.977 |
| RNAInverse | 0.297 | 0.505 |
| MCTS-RNA | 0.667 | 0.793 |
| SAMFEO | 0.770 | 0.969 |
| DesiRNA | 0.907 | 0.996 |
| RIFT-VAE + RNAInverse | 0.803 | 0.985 |
| RIFT-VAE + MCTS-RNA | 0.757 | 0.878 |
| RIFT-VAE + SAMFEO | 0.810 | 0.988 |
| RIFT-VAE + DesiRNA | 0.913 | 0.995 |

Search-only baselines used 3,600 s. Warm-start pipelines used 1,800 s for RIFT-VAE latent optimization followed by 1,800 s for the downstream solver.

Warm-start initialization improved RNAfold-Correct/RNAfold-MCC values for every downstream solver under the matched 3,600-s total budget. RNAInverse showed the largest improvement, from 0.297/0.505 to 0.803/0.985. MCTS-RNA increased from 0.667/0.793 to 0.757/0.878, SAMFEO from 0.770/0.969 to 0.810/0.988, and DesiRNA from 0.907/0.996 to 0.913/0.995. The small change for DesiRNA is consistent with replica-exchange sampling reducing dependence on the initial sequence.

The equal-total-budget design demonstrates that combining the pretrained proposal distribution with downstream search can outperform the corresponding search-only pipeline in this setting.

### Evaluation on the RNAcentral-derived test set

On the RNAcentral-derived test set, RIFT-VAE achieved a Correct value of 0.833 and an MCC of 0.9941 (Table 4). DesiRNA achieved the highest structural accuracy (0.976/0.9997), whereas SAMFEO and MCTS-RNA also produced MCC values close to 1.0. The larger spread in Correct than in MCC indicates that small residual differences involving only a few nucleotides or base pairs can prevent exact equality even when the global base-pair pattern is highly similar.

**Table 4.** Comparison on the RNAcentral-derived test set under a 10-s per-target budget.

| Model | RNAfold-Correct | RNAfold-MCC |
| --- | --- | --- |
| RIFT-VAE | 0.833 | 0.9941 |
| RNAInverse | 0.251 | 0.5607 |
| MCTS-RNA | 0.601 | 0.9886 |
| SAMFEO | 0.758 | 0.9979 |
| DesiRNA | 0.976 | 0.9997 |

The comparison also illustrates complementary operating points. DesiRNA is preferable when maximizing RNAfold structural exactness is the only objective. RIFT-VAE offers a learned proposal distribution that can incorporate additional sequence-level conditions and may be useful as a standalone generator or warm-start component.

### Family-level covariance-model consistency

We next evaluated whether designed sequences retained family-level sequence-structure signatures beyond RNAfold agreement. Under Infernal cmscan with family-specific GA thresholds, the tested search-based designs, including DesiRNA, did not yield correct-family hits in this analysis. RIFT-VAE generated sequences with measurable correct-family annotation rates across multiple RNA families (Figure 6).

**Figure 6.**
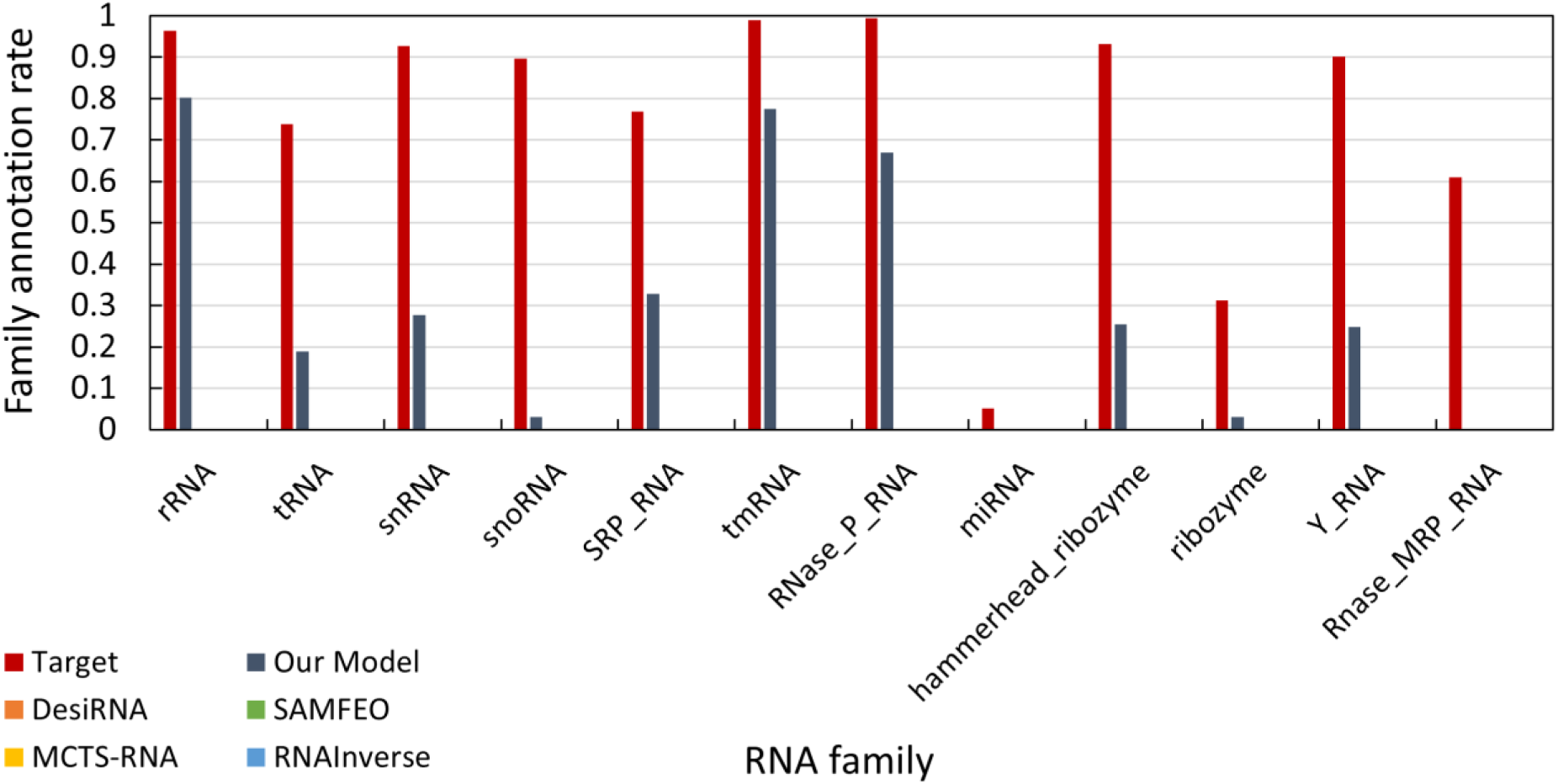
Correct-family annotation rates on the RNAcentral-derived test set. Annotation was performed with Infernal cmscan using family-specific GA thresholds. A positive hit indicates compatibility with a covariance-model family signature but does not by itself establish biological function; absence of a hit does not prove non-functionality.

This difference is consistent with the role of large-scale pretraining: the decoder is trained on natural sequences and can therefore retain conserved combinations of sequence and structure that are not included in a folding-only objective.

### GC-content-controlled generation

Providing a target GC fraction as an explicit input condition improved agreement between generated and specified GC content across the RNAcentral families (Figure 7). MCTS-RNA also accepts a GC-content condition, and its performance reflects this capability. Methods without an explicit GC objective showed larger deviations for several families.

**Figure 7.**
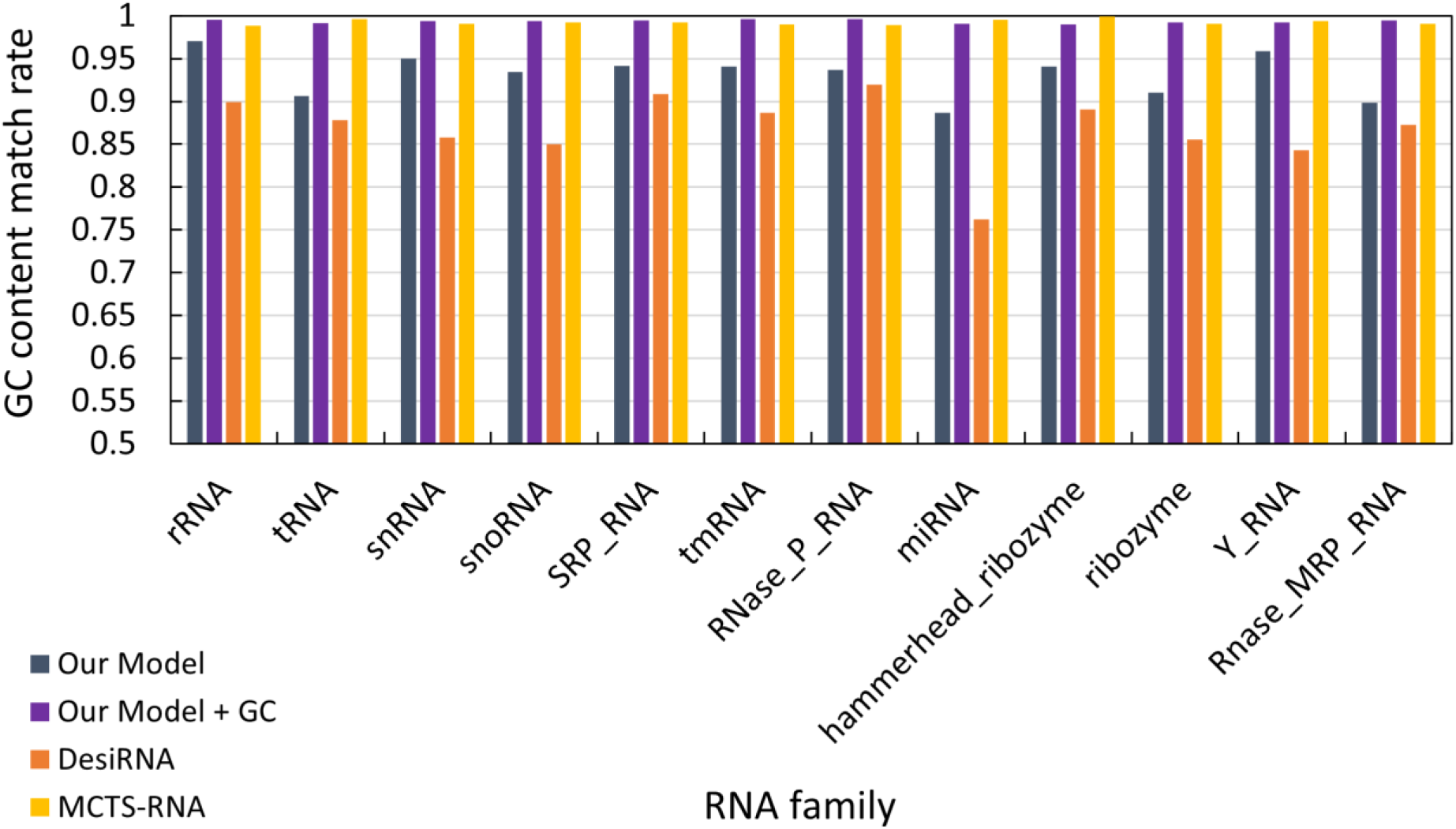
GC-content match across RNA families on the RNAcentral-derived test set. The score is 1 − |g − t|, where g and t denote the generated and target GC fractions, respectively. The GC-conditioned RIFT-VAE variant more closely follows specified targets than its unconditioned counterpart, while MCTS-RNA also benefits from an explicit GC-content condition.

GC content is a simple example of multi-constraint design. The same conditioning interface could be extended to motif requirements, prohibited subsequences, accessibility, energy windows, or application-specific compositional constraints, provided that appropriate training labels and evaluation criteria are available.

### Case study: sequence generation for a hammerhead ribozyme

A type I hammerhead ribozyme was used as a qualitative interpretability case study (Figure 8). The catalytic core and cleavage-site region are strongly conserved in this family. The RIFT-VAE design reproduced the nucleotides corresponding to the catalytic core and matched many conserved positions near the core; it also produced a correct-family covariance-model hit. The DesiRNA design reproduced fewer of these conserved positions and did not yield a correct-family hit under the same analysis.

**Figure 8.**
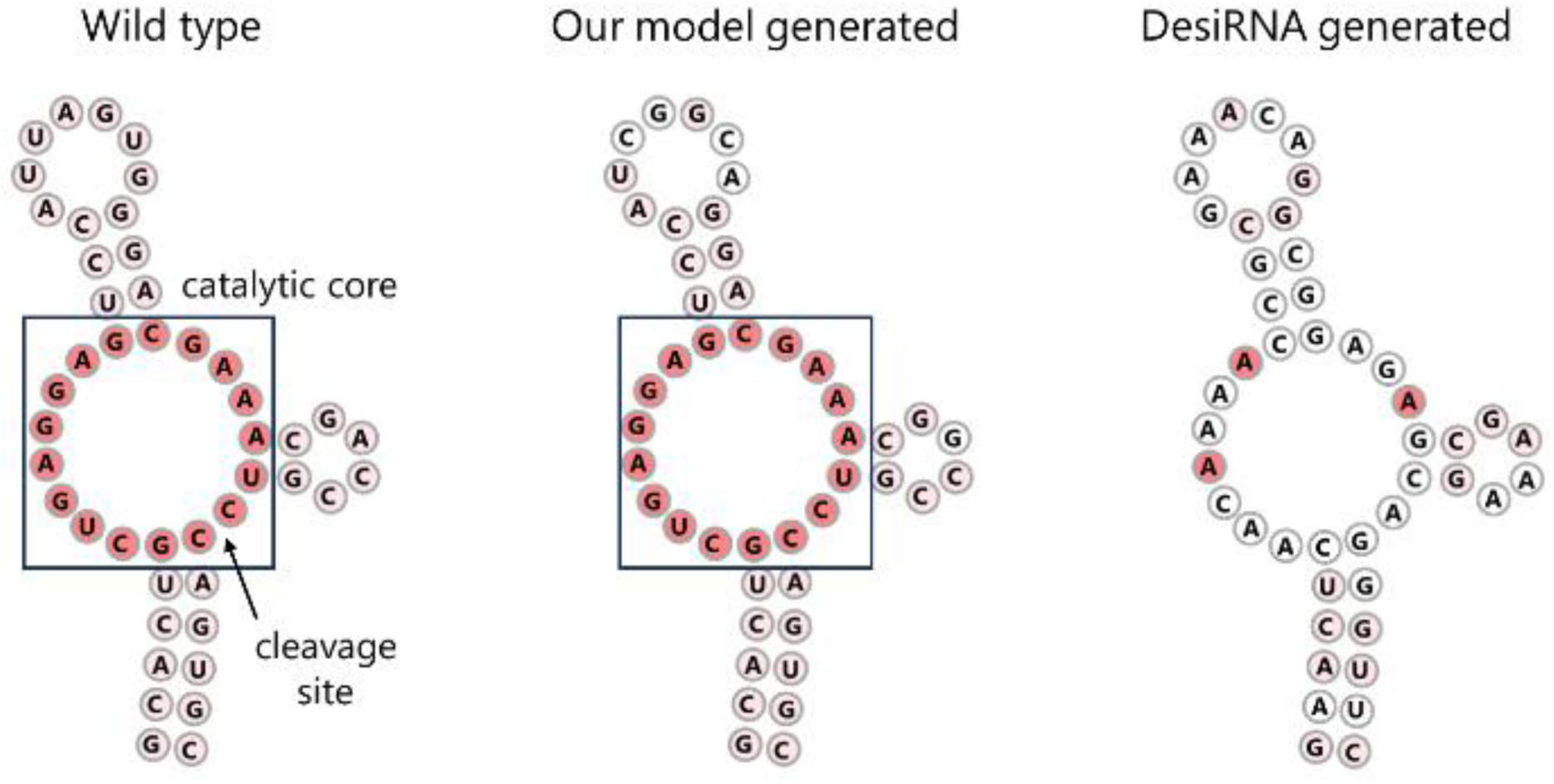
Type I hammerhead ribozyme case study. RNAfold-predicted MFE structures are shown for the reference sequence, the RIFT-VAE design, and the DesiRNA design. Red nucleotide labels identify conserved positions. The RIFT-VAE sequence recapitulates the catalytic-core and cleavage-site region at the sequence level; the figure does not establish catalytic activity or the canonical three-dimensional fold.

All structures shown in Figure 8 are RNAfold-predicted MFE structures. The depicted topology may not faithfully reproduce the canonical three-dimensional hammerhead fold, and activity can depend on long-range loop-loop contacts, metal ions, and tertiary organization. The example therefore demonstrates sequence-level motif recovery under the model, not catalytic activity. Experimental cleavage assays would be required for functional confirmation.

## Discussion

### Performance improvement through four-step enhancements

This study proposes a grammar-conditioned CVAE framework for RNA inverse folding. This work clarifies the role of the CFG parse tree, the transformation from dot-bracket notation to production-rule tokens, and the reconstruction of tree-labeled nucleotides into linear sequences. The key practical finding is that a pretrained generative model can serve both as a standalone sequence generator and as an initializer for search-based solvers.

The performance progression also clarifies the source of improvements. The deterministic Transformer provides a baseline mapping from structure tokens to nucleotides. The CVAE better models the one-to-many nature of inverse folding. Grammar expansion makes local structural context more explicit. SRL reduces inconsistent generations by learning from failures. Latent-space optimization then searches the learned latent space for rare high-agreement candidates. Because the final step contributes the largest gain, future work should compare the same cross-entropy optimization applied to simpler baselines, including dot-bracket-conditioned latent models, to isolate how much benefit comes specifically from the grammar-conditioned CVAE representation.

In sequence design with our proposed model, MCC, which reflects secondary-structure similarity, was close to 1.0, whereas Correct, which reflects the exact-match rate, remained around 0.8. This indicates the difficulty of making fine adjustments to only a few nucleotides or base pairs. Further improvement in Correct will require a more rigorous latent space for nucleotide sequences conditioned on secondary structure, where local movements in latent space do not cause excessively discontinuous changes in generated sequences and their MCC values.

### A learned proposal distribution combined with explicit search

The final performance should be attributed to the complete pipeline rather than to the neural generator alone. The CVAE and pretraining define a conditional proposal distribution over sequences, grammar expansion changes the conditioning representation, SRL reshapes the distribution using predictor-consistent generated pairs, and CEM devotes computation to high-scoring latent regions. On EteRNA100, the jump from 0.250 to 0.760 Correct after latent optimization shows that explicit search is the dominant final step.

### Practical positioning relative to search-based solvers

The results suggest three use cases. First, when exact RNAfold-MFE recovery is the only objective, DesiRNA provides the strongest performance among the evaluated methods. Second, when designs should also preserve family-level covariance-model signatures or satisfy an explicit GC condition, RIFT-VAE provides an integrated learned interface. Third, when an established local or heuristic solver is available, RIFT-VAE can supply a warm start that improves the equal-budget pipeline, particularly for methods sensitive to initialization.

This positioning is more informative than claiming universal superiority. Different applications place different weights on exact folding-model agreement, sequence naturalness, diversity, runtime, additional constraints, and experimental manufacturability. A design framework should therefore expose these trade-offs rather than reduce them to a single score.

### Beyond structural correctness: family features and design constraints

The family annotation analysis highlights a complementary objective to structural correctness. Search-based methods optimize the specified structural or energy score and may produce structurally valid sequences that lack natural family signatures. Our model, trained on large-scale naturally derived sequences, produces some designs that retain covariance-model family signals. However, a family annotation hit is only a proxy for sequence/structure compatibility with a known RNA family, and failure to hit a covariance model does not prove non-functionality. Practical applications should identify specific families where conserved motifs are known to be functionally required and should validate designs experimentally.

For GC content control, we showed that by providing a specified GC content value as an input condition, the model can generate sequences consistent with the target GC content. In applications such as temperature-responsive RNAs, where GC content can correlate with function, the ability to incorporate sequence-level properties such as GC content as conditions alongside secondary structure is a practical advantage.

### Limitations and validation priorities

A central limitation is the reliance on RNAfold. RNAfold is used to generate secondary-structure labels for RNAcentral, to validate generated sequences, and to construct SRL examples. Consequently, the reported Correct and MCC scores measure agreement with the RNAfold MFE model rather than agreement with experimentally determined structures. This does not invalidate the benchmark comparison, because many inverse-folding benchmarks use RNAfold-based validation, but it limits biological interpretation. Orthogonal validation with independent predictors, such as a classical physics-based tool like RNAstructure and a recent machine-learning-based predictor such as BPfold, would be necessary to reduce the risk that the model is merely learning to invert the RNAfold energy model [20,21].

The model is also restricted to pseudoknot-free secondary structures and canonical base-pairing information represented by dot-bracket notation. Functional RNAs can depend on tertiary contacts, pseudoknots, triplexes, kissing loops, and non-Watson-Crick base pairs. The CFG representation can in principle be extended by adding labels for non-canonical base-pair types, pseudoknot modules, or graph-based tertiary-contact constraints, but such information is not included in the present model. Therefore, our outputs should be considered candidates for further computational and experimental screening rather than validated functional RNAs.

Finally, our current evaluation reports the best-scoring candidate among generated samples. This is appropriate for design success but does not fully characterize diversity. CVAE sampling is motivated partly by the ability to generate multiple valid sequences for the same structure, whereas SRL and latent-space optimization can reduce diversity by focusing on high-scoring regions. Future evaluations should report sequence diversity, for example pairwise normalized Hamming distance among candidates at fixed MCC thresholds, and should analyze the trade-off between diversity and exact structural recovery.

The family-holdout analysis should also be interpreted cautiously. Although scaRNA, scRNA, and telomerase RNA families were excluded from training, distantly related families from the same broader clans may remain in the training set. A more stringent future test would exclude entire clans or use additional sequence- and structure-based de-duplication. In addition, our current evaluation reports best-of-sample design success and does not fully quantify diversity among generated sequences; future work should report diversity at fixed structural quality.

## Conclusion

RIFT-VAE provides a grammar-conditioned, pretrained framework for RNA inverse folding that combines conditional generation with explicit latent-space search. The CFG parse tree offers a structured interface for stem-loop hierarchy and base-pair context, while the CVAE, SRL, and CEM components progressively improve compatibility with target structures under RNAfold. The method also supports GC-content conditioning, retains measurable family-level covariance-model signatures, and supplies useful initial solutions to search-based solvers.

RIFT-VAE provides a useful bridge between classical RNA secondary-structure representations and modern generative sequence design. The method is strongest when high secondary-structure agreement is desired together with natural-like sequence features or when a good initialization is needed for downstream search. The next steps are orthogonal validation, stricter redundancy-controlled data splits, diversity-aware evaluation, and extension beyond RNAfold-predicted pseudoknot-free secondary structures.

## Acknowledgements

We thank the members of the Sakakibara laboratory for helpful discussions and feedback on this work.

## Author contributions

K.W. implemented the model, conducted the experiments, and analyzed the data. M.A. contributed to methodology, analysis, and interpretation of the results. Y.S. conceived and supervised the study, secured funding, interpreted the results, and drafted and revised the manuscript. All authors read and approved the final manuscript.

## Data availability

The RNAcentral-derived data splits, processed datasets, and pretrained model resources supporting this study are available through Zenodo at https://doi.org/10.5281/zenodo.18678647 and are cited as a dataset in the reference list [22]. RNAcentral and EteRNA100 are third-party resources available from their original providers [10, 14].

## Code availability

Source code for preprocessing, model training, sequence generation, SRL, latent-space optimization, and benchmark evaluation is available at https://github.com/Kentaro0727/RIFT-VAE. The repository includes the grammar definitions and an environment specification.

## References

1. Jumper J, Evans R, Pritzel A, et al. Highly accurate protein structure prediction with AlphaFold. Nature 2021; 596:583–589.

2. Abramson J, Adler J, Dunger J, et al. Accurate structure prediction of biomolecular interactions with AlphaFold 3. Nature 2024; 630:493–500.

3. Watson JL, Juergens D, Bennett NR, et al. De novo design of protein structure and function with RFdiffusion. Nature 2023; 620:1089–1100.

4. Wong F, et al. Deep generative design of RNA aptamers using structural predictions. Nat Comput Sci 2024; 4:829–839.

5. Hofacker IL, Fontana W, Stadler PF, Bonhoeffer LS, Tacker M, Schuster P. Fast folding and comparison of RNA secondary structures. Monatsh Chem 1994; 125:167–188.

6. Lorenz R, Bernhart SH, Hoener zu Siederdissen C, et al. ViennaRNA Package 2.0. Algorithms Mol Biol 2011; 6:26.

7. Yang X, Yoshizoe K, Taneda A. RNA inverse folding using Monte Carlo tree search. BMC Bioinformatics 2017; 18:468.

8. Zhou T, et al. RNA design via structure-aware multifrontier ensemble optimization. Bioinformatics 2023; 39:i563–i571.

9. Wirecki TK, et al. DesiRNA: structure-based design of RNA sequences with a replica exchange Monte Carlo approach. Nucleic Acids Res 2025; 53:gkae1306.

10. Koodli RV, Rudolfs B, Wayment-Steele HK, Das R. Redesigning the EteRNA100 for the Vienna 2 folding engine. bioRxiv 2021; 2021.08.26.457839.

11. Patil S, et al. RNAinformer: generative RNA design with tertiary interactions. bioRxiv 2024.

12. Huang H, et al. RiboDiffusion: tertiary structure-based RNA inverse folding with generative diffusion models. Bioinformatics 2024; 40:i347–i356.

13. Dotu I, et al. Complete RNA inverse folding: computational design of functional hammerhead ribozymes. Nucleic Acids Res 2014; 42:11752–11762.

14. RNAcentral Consortium. RNAcentral 2021: secondary structure integration, improved sequence search and new member databases. Nucleic Acids Res 2021; 49:D212–D220.

15. Vaswani A, Shazeer N, Parmar N, et al. Attention is all you need. Adv Neural Inf Process Syst 2017; 30.

16. Sohn K, Lee H, Yan X. Learning structured output representation using deep conditional generative models. Adv Neural Inf Process Syst 2015; 28.

17. Rubinstein R. The cross-entropy method for combinatorial and continuous optimization. Methodol Comput Appl Probab 1999; 1:127–190.

18. de Boer PT, Kroese DP, Mannor S, Rubinstein RY. A tutorial on the cross-entropy method. Ann Oper Res 2005; 134:19–67.

19. Nawrocki EP, Eddy SR. Infernal 1.1: 100-fold faster RNA homology searches. Bioinformatics 2013; 29:2933–2935.

20. Reuter JS, Mathews DH. RNAstructure: software for RNA secondary structure prediction and analysis. BMC Bioinformatics 2010; 11:129.

21. Zhu H, et al. Deep generalizable prediction of RNA secondary structure via base pair motif energy. Nat Commun 2025; 16:5856.

22. [dataset] Watanabe K, Akiyama M, Sakakibara Y. A pretrained model for RNA inverse folding with formal grammar representations. Zenodo, 2026. 10.5281/zenodo.18678647.

